# MANGIFERIN AS A POTENTIAL INHIBITOR OF TRANSTHYRETIN FIBRILLOGENESIS

**DOI:** 10.1101/2024.10.09.617482

**Authors:** N.A. Grudinina, M.G. Petukhov, A.A. Shaldzhyan, Y.A. Zabrodskaya, N.V. Gavrilova, S.N. Morozkina, V.V. Egorov

**Affiliations:** Federal State Budgetary Scientific Institution “Institute of Experimental Medicine”, 197022, Akademika Pavlova street 12, St. Petersburg, Russia; Petersburg Nuclear Physics Institute named by B.P. Konstantinov of National Research Centre “Kurchatov Institute”, 188300, mkr. Orlova roshcha 1, Gatchina, Russia; Smorodintsev Research Institute of Influenza, Russian Ministry of Health, 197376, Prof. Popov St. 15/17, St. Petersburg, Russia; Institute of Biomedical Systems and Biotechnology, Peter the Great Saint-Petersburg Polytechnic University, 194064, Politekhnicheskaya 29, St. Petersburg, Russia; ITMO University, 197101, Kronverksky Pr. 49, bldg. A, St. Petersburg, Russia; Kabardino-Balkarian State University named after H.M. Berbekov, 360004, Chernyshevskogo street, 173, Nalchik, Kabardino-Balkaria, Russia; Federal State Budgetary Educational Institution of Higher Professional Education “Saint-Petersburg State University”, 199034, Universitetskaya embankment, 7-9, St. Petersburg, Russia

**Keywords:** TRANSTHYRETIN, AMYLOIDOSIS, FLUORIMETRY, RECOMBINANT PROTEINS, ANTIAMYLOIDOGENIC DRUGS, MOLECULAR DOCKING

## Abstract

By using *in silico* molecular docking, we predicted a high potential of mangiferin to inhibit transthyretin amyloidogenesis. Subsequently, we tested this prediction in an *in vitro* system. The study involved testing effect of mangiferrin in a system that included isolated and purified recombinant transthyretin under fibrillogenesis conditions and a detection assay based on the fluorescent dye thioflavin T, characteristic of amyloid fibrils. The conditions for fibrillogenesis of this protein at neutral pH values were established. Using the developed test system, it was shown that mangiferin is able to inhibit abnormal transthyretin fibrillogenesis.

## Introduction

Transthyretin (TTR) is a serum and cerebrospinal fluid protein that transports retinol-binding protein and thyroxine. It is a homotetrameric protein synthesized primarily in the liver, choroid plexus, retinal pigment epithelium, and pancreas. TTR is capable of forming amyloid-like fibrils. Fibrillogenesis of TTR in the body leads to the formation of extracellular protein deposits that impair organ function. Transthyretin oligomers are also capable of inducing cell apoptosis [1]. Clinical syndromes associated with TTR aggregation include familial amyloid polyneuropathy (FAP) and cardiomyopathy (FAP), in which mutant TTR protein aggregates in the peripheral and autonomic nerves and the heart, respectively; and senile systemic amyloidosis (SSA), in which the wild-type protein is deposited primarily in the heart, as well as in the intestine and carpal tunnel. Both wild-type transthyretin and transthyretin carrying amino acid substitutions are capable of fibrillogenesis. Currently, more than 50 mutations in the human transthyretin gene are known, most of them lead to the development of transthyretin amyloidosis (ATTR). Some of the amino acid substitutions leading to the development of ATTR cause a decrease in the stability of the native tetramer. As has been shown, dissociation of the TTR tetramer is the first step leading to the formation of non-native quaternary structures and the formation of amyloid fibrils [2]. Therefore, one of the directions of searching for therapeutic agents to combat ATTR is the search for low-molecular compounds that stabilize the tetramer. [3].

A characteristic feature of amyloid fibrils is the ability to form complexes with the dye thioflavin T and increase the quantum yield of its fluorescence, as well as to shift the absorption spectrum of the dye Congo red to the long-wave region. [4]. Changes in the spectral properties of these dyes in complexes with amyloid fibrils are used in the diagnosis of amyloidosis. [5] and in conducting studies of amyloid-like fibrillogenesis *in vitro*. The increase in the quantum yield of fluorescence of thioflavin T in complex with amyloid-like fibrils (with excitation at a wavelength of 440 nm and detection at 478-482 nm) is described in the basic work [6].

Mangiferin (2-beta-D-glucopyranosyl-1,3,6,7-tetrahydroxy-9H-xanthen-9-one, MW=422.3 g/mol) a substance isolated from mangoes that has no proven therapeutic effect but having a large number of potential pharmacophores [7]. Preliminary *in vitro* screening has shown that xanthans, in particular mangiferrin, may have a modulating effect on TTR oligomerization. In this study, we test the effect of mangiferin on transthyretin fibrillogenesis *in vitro*.

## Materials and methods

### 1.1 Materials

All reagents and materials used in this study were purchased from Sigma (USA) and Merck (USA).

### 1.2 Molecular docking

A full-atom model of the spatial structure of amyloid fibrils of the TTR were constructed based on the crystal structure from the Protein Data Bank (PDB ID: 6SDZ) [8]. The missing loop in the crystal structure (amino acid residues 36-56) was reconstructed using standard protocols in ICM-Pro (Molsoft LLC, USA) [9] with calculations utilizing the ECEPP/3 force field parameters implemented in the ICM-Pro software package as described in [10]. The spatial structure of the mangiferin was taken from PubChem database. The standard algorithm of virtual screening of flexible ligands from the ICM-Pro software package was used for mangiferin docking in the 3 internal cavities available in the spatial structure of TTR amyloid fibrils. The search for the lowest-energy conformations of the ligand in a binding pocket of TTR amyloid fibrils and additional relaxation of ligand steric stains were done as described in [10].

### 1.3 Genetic construct

We used a genetic construct based on the expression plasmid pET22b+ from the collection of the Department of Molecular Genetics of the Institute of Experimental Medicine, obtained from the construct described in [11], but carrying a nucleotide substitution resulting in the amino acid substitution D38H (pet-TN). The nucleotide sequence of the insert and the corresponding amino acid sequence of the recombinant protein are given in Supplement 1. Sequencing was carried out by employees of the resource center of the Federal State Budgetary Scientific Institution IEM.

### 1.4 Isolation and purification of recombinant protein

The recombinant TTR expressed in the *E. coli* BL21DE3 producent was isolated similarly to the procedure described in [11]. Briefly, the bacterial medium containing the recombinant TTR was diluted 2-fold with distilled water and loaded onto a 35 ml column with DEAE A50 Sephadex. Loading was carried out under pressure for 17 hours, then the column was washed with 300 ml PBS and the protein was removed with a step gradient of NaCl (200-250-300-350 mM). Protein elution was monitored by absorption at 280 nm using a NanoDrop spectrophotometer (Thermo, USA). The fractions obtained by ion exchange chromatography were then concentrated using an Amicon concentrator (10 kDa filter), followed by gel filtration on a HiLoad 26/60 Superdex 200 pg column. The mobile phase was phosphate buffered saline, pH 7.2; flow rate was 2.6 ml/min (29 cm/h). The fraction containing the main peak of the transthyretin tetramer was again concentrated on an Amicon (10 kDa).

### 1.5 Polyacrylamide gel electrophoresis (PAGE)

Denaturing electrophoresis was performed according to the method [12]. 5 μl of a solution containing 2% SDS, 2.5% beta-mercaptoethanol and 0.01% bromophenol blue were added to 5 μl of protein at a concentration of about 1 mg/ml, then some of the samples were incubated in a boiling water bath for 3 minutes, while others were applied without such incubation. Protein electrophoresis in 12% PAAG was performed in 10×10 cm plates at a voltage gradient of 20 V/cm. The gel was stained with Coomassie R-250 solution according to [13]; images of the stained gel were obtained in a ChemiDoc MP gel-documenting station (BioRad, USA).

### 1.6 Mass spectrometry analysis

To analyze the amino acid sequence of the protein after PAGE, enzymatic hydrolysis in the gel with trypsin was performed. A fragment of the stained zone was cut out, washed from the dye (twice with 100 μl of a solution of 30 mM ammonium bicarbonate, 40% acetonitrile in water), dehydrated in 100% acetonitrile, after which acetonitrile was collected and the gel fragment was incubated in air until complete evaporation of acetonitrile. Then 2 μl of trypsin solution (20 μg/ml in 50 mM ammonium bicarbonate) were added to the gel fragment and incubated in a solid-state thermostat GNOM (“DNK-Tecknologii”, Russia) at 37°C for 18 hours. The reaction was stopped with 3 μl of a solution of 1% TFA, 10% acetonitrile in water. To identify proteins, the resulting set of tryptic peptides was mixed with the DHB matrix in equal volumes, applied to a steel target and analyzed in the reflective mode of positive ion registration on an UltrafleXtreme (Bruker, Germany) MALDI-TOF/TOF mass spectrometer. At least 5000 laser pulses were summed for each spectrum.

Protein identification was performed using MASCOT [14] by simultaneously accessing the SwissProt database [15]and the local database, where the sequence of the transthyretin recombinant protein (called “TTR_rec”) was previously added. The accuracy of mass determination was limited to 50 ppm. Up to two trypsin errors (missing a proteolytic site) were allowed.

### 1.7 Transmission Electron Microscopy (TEM)

For TEM, 5 μl of 1 mg/mL TTR fibrils suspension was applied to copper grid for electron microscopy coated with a carbon film. After adsorption of the proteins to the grid surface for 1 min, then the grids were washed twice with distilled water. Next, negative staining of the samples on the grids was performed with a 2% solution of sodium phosphotungstic acid salt for 1 min. After staining, the grids were dried and examined in a JEOL JEM 1011 transmission electron microscope at an accelerating voltage of 80 kV. Electron micrographs were obtained using a Morada digital camera (Olympus Inc.)

### 1.8 Measurement of thioflavin T fluorescence

20 μl of the sample were added to 80 μl of the buffer containing thioflavin T (to a concentration of 10 μM thioflavin T), thoroughly mixed and incubated for 15 minutes in the dark. Measurements were performed in 96-well darkened NUNC plates using a BMG Clariostar plate spectrofluorimeter in fluorescence mode with an excitation wavelength of 440-10 nm, fluorescence was recorded at 478-10 nm. The signal gain value was set based on the fluorescence maximum in the studied series of samples. The signal was recorded along the well bottom using integration over the entire area of the cell. The results of at least three measurements were used for processing. The SPSS SigmaPlot program was used to process and visualize the results. Statistical data analysis was performed in the GraphPad PRISM 9.0 program using ordinary one-way ANOVA (correction for multiple comparitions were performed using Sidak test): ^****^—p < 0.0001; ^***^—p < 0.001

## 2. Results

### 2.1 Molecular docking

Figure 1 shows the molecular model used for molecular docking of mangiferin in the inner cavity of TTR amyloid fibrils containing 2, 3 and 5 chains of the transthyretin protein.

**Figure 1.**
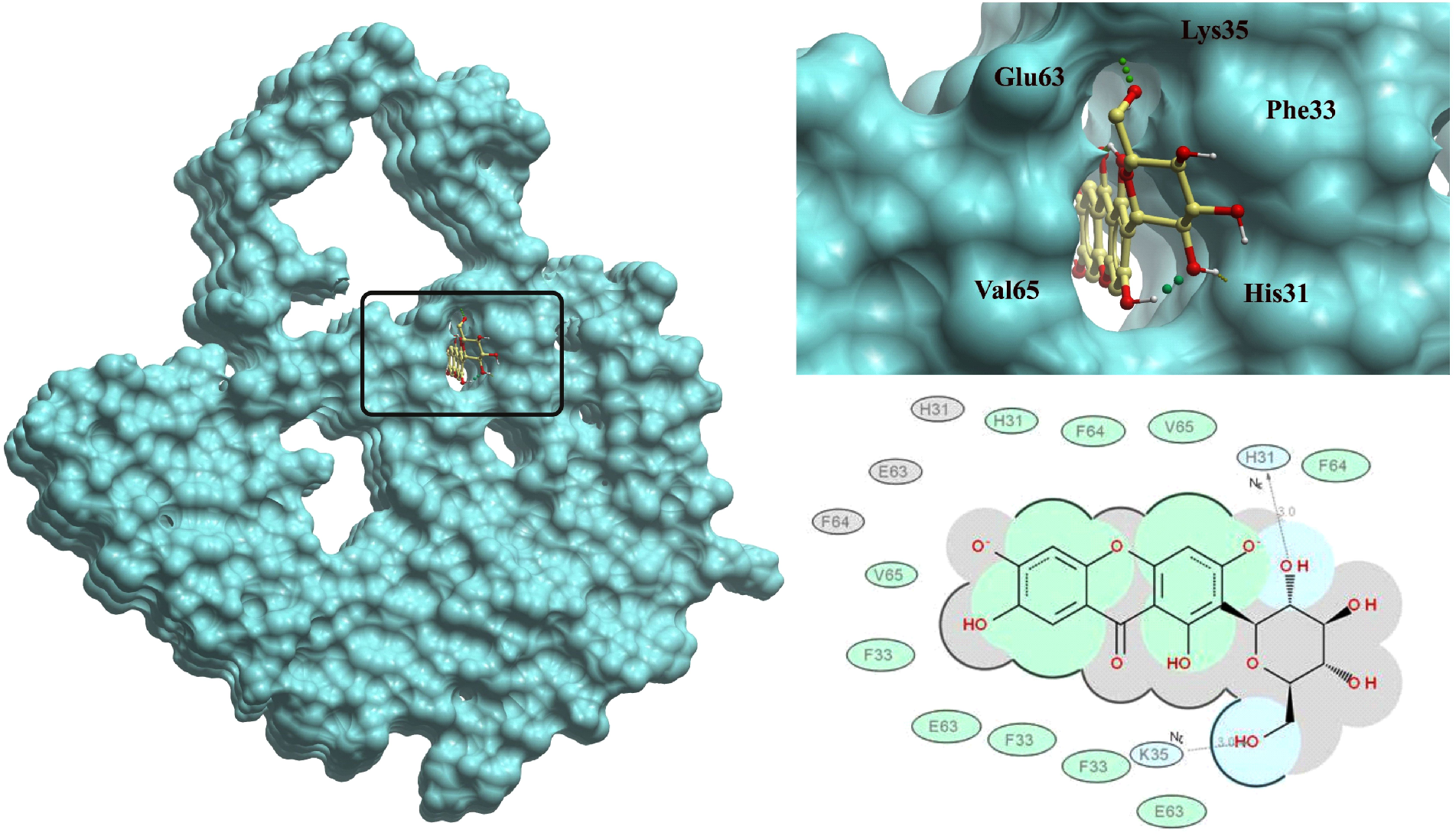
Spatial structure of the TTR amyloid fibrils in the complex with the drug-like organic compound mangiferin in its internal cavity obtained using VLS the calculations.

In this work, we compared the ICM-score values obtained using the docking algorithm for binding mangiferin in three cavities observed in the TTR amyloid fibril. The best docking score (ICM score=-24.5) was obtained for the internal cavities shown in Figure 1. It was found that the other two cavities were either too large or too small for effective binding of mangiferin. The structure of the complex shown in Figure 1 is characterized by a large buried hydrophobic ligand surface located in the central part of the cavity. There are also two hydrogen bonds between the terminal hydroxyl groups of the ligand and His31 and Lys35 of the TTR protein. Other hydroxyl groups of the ligand either form intramolecular hydrogen bonds or are well hydrated by the surrounding solvent. Interestingly, the ICM-Score values also depend on the size of the amyloid fibrils, while the minimum ICM-Score is observed for amyloid fibrils containing three TTR protein chains. A further increase in the size of the amyloid fibril worsens the hydrogenation conditions of the hydroxyl groups of the ligand at one end of the inner cavity of the amyloid fibril.

### 2.2 Production of recombinant transthyretin

IPTG-induced expression of the transthyretin gene from the pet-TN plasmid was performed in the *E. Coli* expression strain BL21DE3 using standard methods [16]. The protein was then isolated from the culture medium using ion-exchange chromatography on DEAE-Sepharose. The fractions obtained were analyzed using PAGE. Fractions containing the target protein were concentrated and sent for purification by gel filtration (see Materials and Methods). Figure 2 shows the electropherogram of the purified TTR obtained.

**Figure 2.**
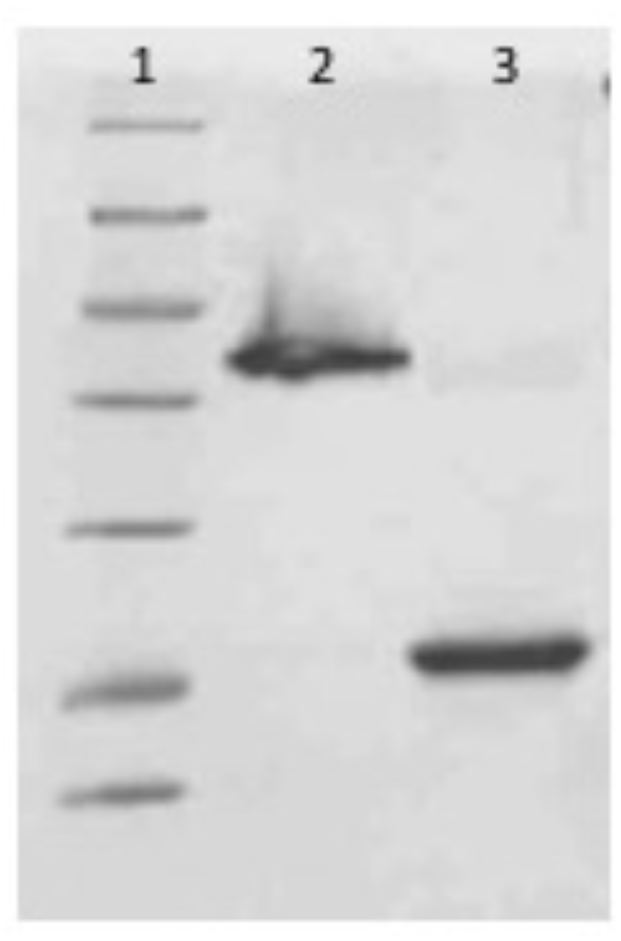
Electrophoregram of purified transthyretin. Lane 1 shows the molecular weight marker (Thermo Scientific Unstained Protein Molecular Weight Marker 26610), lane 2 shows the TTR sample that was not subjected to heat denaturation, and lane 3 shows the TTR sample after boiling.

The protein purity was over 95%, and the obtained bands were used for mass spectrometric analysis. The obtained mass spectra (see Supplement) were used to reliably identify the recombinant protein TTR_rec (transthyretin recombinant protein); the Score value was 228 with a threshold value of 70 (if the Score exceeds the threshold value, the identification is considered reliable, p < 0.05). The protein sequence coverage (the percentage of amino acid residues to which the ions from the spectrum were matched) was 84%. Thus, we confirmed the primary structure of the isolated recombinant protein.

### 2.2 Obtaining of transthyretin fibrils at neutral pH

The obtained protein was used to select the conditions for fibrillogenesis. According to the study described in [17], low pH values and the presence of EDTA are optimal for fibrillogenesis of this protein. The interaction of the potential drug with the protein should occur under conditions close to physiological, therefore, the *in silico* modeling of the interaction of the studied compound with transthyretin was carried out by simulating pH close to physiological (pH=7). To verify the modeling results in *in vitro* experiments, a search for conditions for transthyretin fibrillogenesis at neutral pH values was carried out. Fibril formation was detected by a change in thioflavin T fluorescence as described in Materials and Methods section. Varying the NaCl concentration made it possible to find conditions for fibrillogenesis at pH 7. The following conditions were used for fibrillogenesis of the obtained transthyretin: Transthyretin concentration - 1 mg / ml, NaCl - 200 mM, phosphate buffer pH 7.2 -9 mM, GlyHCl buffer pH 2 - 5 mM, EDTA - 50 μM; A 100 µl sample was incubated at 37°C for 24 hours without stirring. Electron microscopy showed the presence of aggregates in the studied sample, morphologically corresponding to amyloid-like fibrils. Figure 3 shows images of fibrils obtained at neutral pH (in the presence of 3% DMSO) and the absorption spectrum of Congo Red in the presence of fibrils. [4]. Data on the fluorescence intensity of Thioflavin T in the studied samples are presented in Figure 4 (chart column «Fibrils DMSO»).

**Figure 3.**
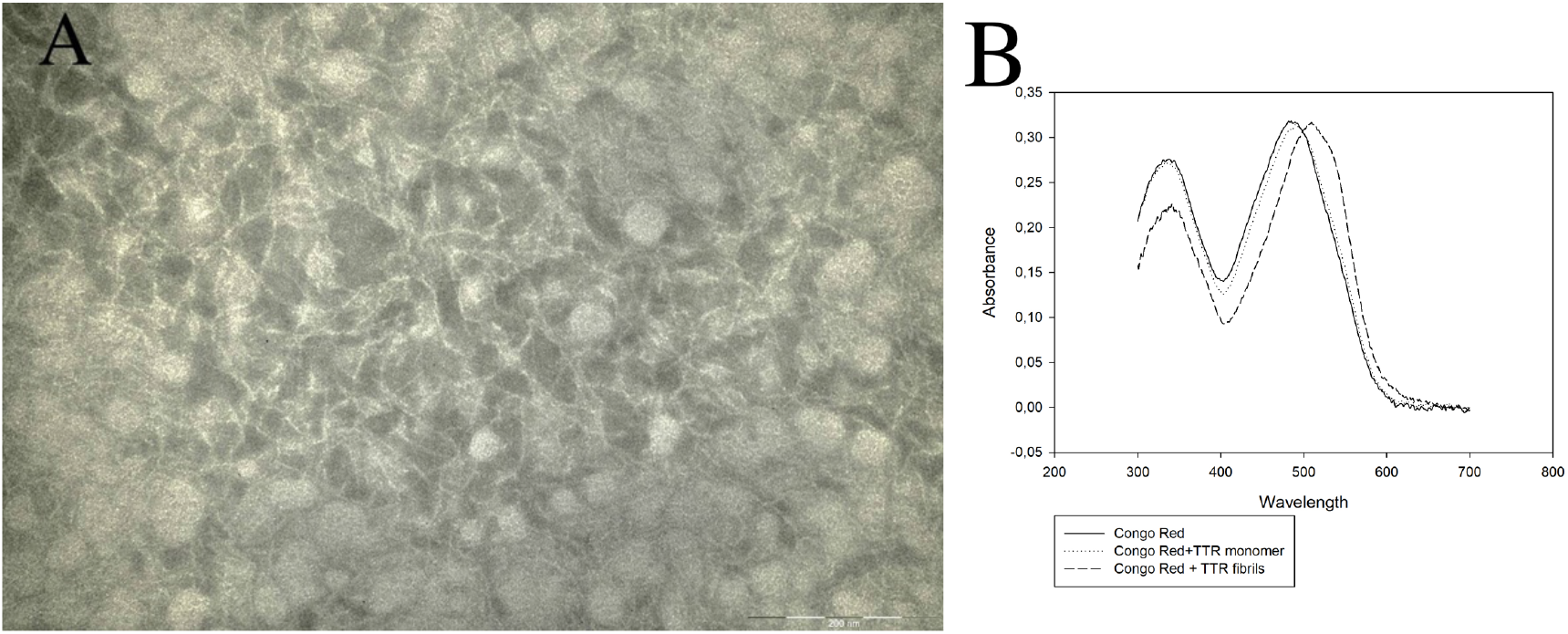
(A) Electron microscopic images of fibrils. The length of the measuring segment is 500 nm in the image on the left and 200 nm and (B) absorption spectrum of CR in the presence of TTR monomers and fibrils.

**Figure 4.**
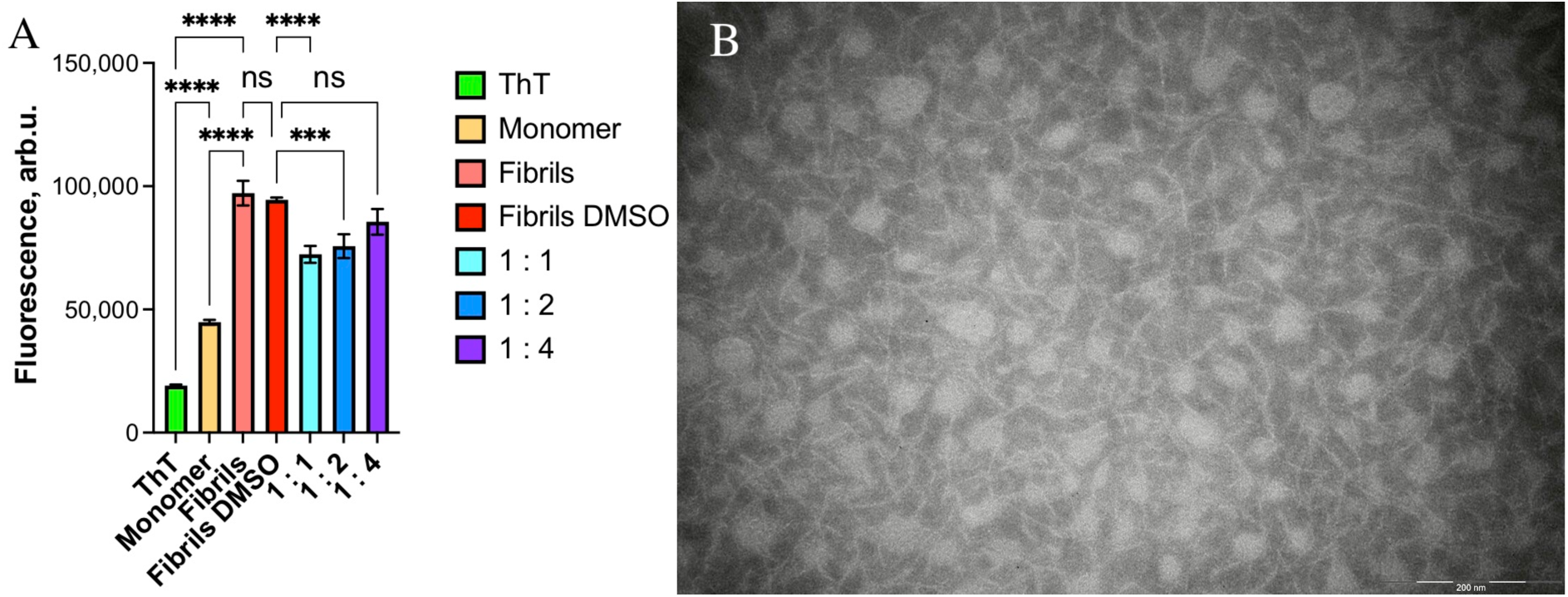
Results of testing mangiferin for antiamyloidogenic activity (A) «ThT» diagram column corresponds to the intensity of Thioflavin T background fluorescence in the fibrillogenesis buffer, «Monomer» in the presence of the monomer, «Fibrils» - in the presence of control transthyretin fibrils, «Fibrils DMSO» - in the presence of control transthyretin fibrils obtained in the buffer with 3% DMSO, «1:1», «1:2», «1:4» corresponds to the fluorescence intensity in the presence of 3% DMSO of fibrils formed in samples containing the corresponding (30 μg/ml (1:1 mangiferin:TTR molar ratio), 15 μg/ml (1:2), and 7.5 μg/ml (1:4) respectively) concentrations of mangiferin; (B) Electron microscopic images of fibrils formed in the presence of mangiferin («1:1» sample). The results of statistical processing of the experimental results showed that: (i) there are no reliable differences between the fluorescence intensity of the ThT complex and fibrils that were formed in the presence and absence of DMSO at the concentrations used in the experiments with mangiferin (ii) the fluorescence of the ThT complexes with fibrils formed in the presence of mangiferin at molar ratio of 1:1 and 1:2 (but not 1:4) is significantly different from that of the control fibrils.

The observed fibrillar morphology of the aggregates and the shift in the absorption spectrum of Congo Red and the increase in the fluorescence intensity of thioflavin T indicate the presence of amyloid-like fibrils in the sample.

### 2.3 Mangiferin testing

Mangiferin (1 mg) was dissolved in 1 ml 100% DMSO. To test the ability of the compound to affect fibrillogenesis, 3 μl of the compound solution (m) were added to 97 μl of the protein in the buffer described in section 2.2, as well as 3 μl of the compound diluted 2 (m1) and 4 (m 0.5) times, respectively, with fibrillogenesis buffer containing DMSO, to the 3% v/v final concentration. Measurements were performed in triplicate. Figure 4 shows the results of measuring the fluorescence intensity of Thioflavin T in the presence of mangiferin, as well as control fibrils and transthyretin in soluble form (red). Control experiments showed that DMSO at the concentration used did not affect the fibrillogenesis process. Based on the comparison of the Fibrils and Fibrils DMSO diagram columns, it can be concluded that 3% DMSO also does not affect fibrillogenesis or the fluorescence of Thioflavin T in the presence of fibrils.

Comparing the images of fibrils in Figures 3A and 4B, it can be seen that in the absence of mangiferin, fibrils do not form, or form to a much lesser extent, intertwined protofibril structures. It is possible that a change in the surface of protofibrils, leading to a change in the ability to interfibrillar interaction, also leads to a change in the ability of protofibrils to bind and change the spectral properties of thioflavin T.

Thus, the test results showed that under the experimental conditions mangiferin had, within the framework of the testing methodology, an antiamyloidogenic effect. It should be noted that mangiferin is soluble under conditions close to physiological up to concentrations corresponding to the concentration of transthyretin (70 μM), and has an antiamyloidogenic effect.

## Conclusion

Based on the results of the study, it can be concluded that the study of mangiferin as a potential pharmacological substance for combating ATTR is promising. Further studies of the biodistribution and pharmacokinetics of mangiferin in animal models are needed to resolve the issue of the prospects for using the drug to combat transthyretin amyloidosis.

## Acknowledgments

This study was supported with RSF project no. 21-74-20093 and is partially funded by the Ministry of Science and Higher Education of the Russian Federation as part of World-class Research Center program: Advanced Digital Technologies (contract No. 075-15-2022-311 dated by 20.04.2022) The authors thank Dr Andrey Kajava for his advices and critical comments.

## Author contribution

Egorov V.V. – experiment design, fluorometry, manuscript writing

Grudinina N.A. – protein expression and purification, fluorometry

Petukhov M.G. – molecular modelling

Shaldzhyan A.A. – protein purification by gel-filtration

Zabrodskaya Y.A. – mass-spectrometry

Gavrilova N.V. – transmission electron microscopy

Morozkina S.N. – mangiderin purification

## Genetic construct encoding transthyretin

A genetic construct based on the expression plasmid pet22b+ encoding human transthyretin was used. The results of sequencing the protein-coding portion of the pet-TN plasmid using standard primers are shown in Figure 1.

**Figure 1.**
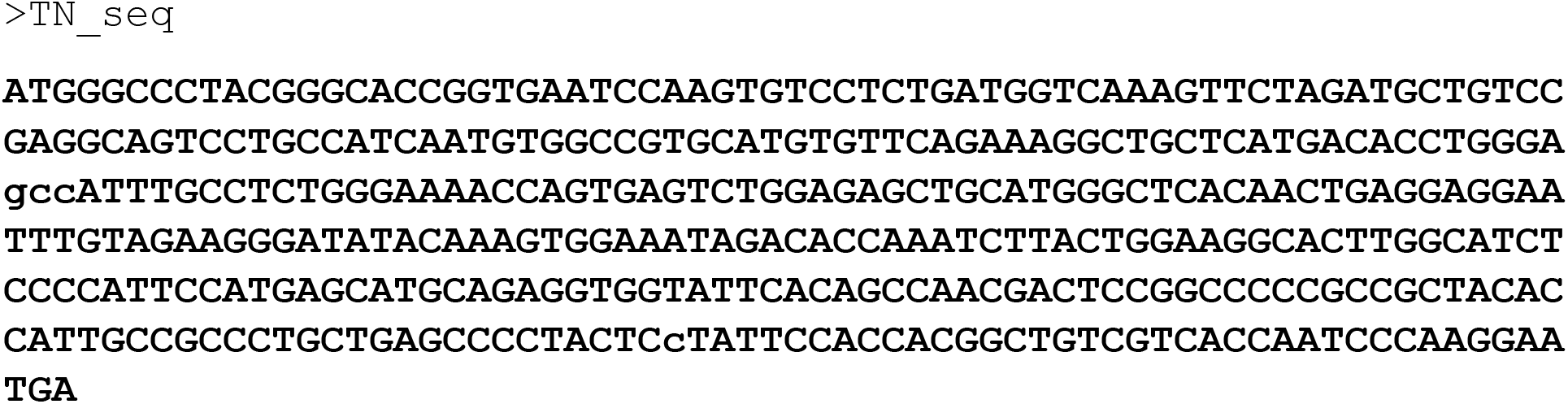
Results of sequencing of a section of the pet22b+ plasmid containing the transthyretin gene. The coding sequence of the recombinant protein is shown in bold.

Figure 2 shows the amino acid sequence of the protein encoded by the sequence inserted into the expression plasmid. The amino acid substitution D38H is shown in bold.

**Figure 2.**
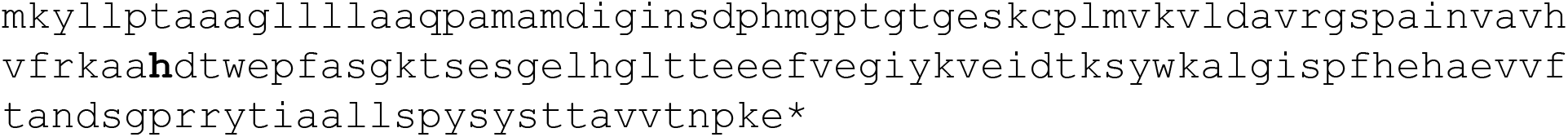
Amino acid sequence of recombinant transthyretin. The amino acid substitution D38H is shown in bold.

**Figure 3:**
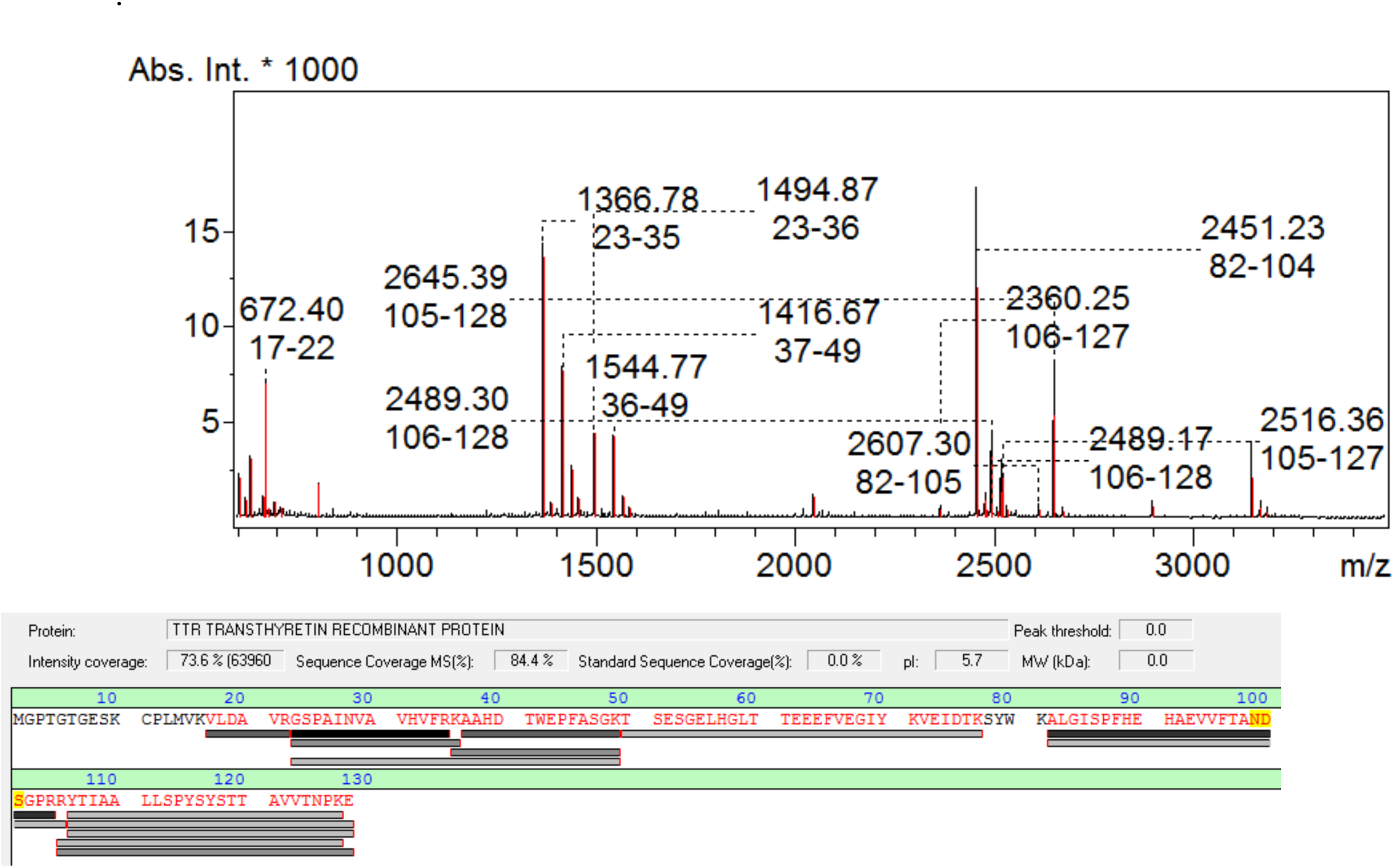
(Top) Fragment of the MALDI mass spectrum obtained after tryptic hydrolysis of the colored PAGE zone. Ions associated with fragments of the amino acid sequence of the TTR recombinant protein and the corresponding range of amino acids are marked. (Bottom) Amino acid sequence of the TTR_rec protein, amino acids detected in the spectrum are marked in red; peptides associated with the registered ions are underlined in gray.

